# Potent photoswitch for expression of biotherapeutics in mammalian cells by light

**DOI:** 10.1101/2024.10.03.616529

**Authors:** Jeannette Gebel, Elisa Ciglieri, Rainer Stahl, Fraser Duthie, Andreas Möglich, Herbert Müller-Hartmann, Hanns-Martin Schmidt, Dagmar Wachten

## Abstract

Precise temporal and spatial control of gene expression is of great benefit for the study of specific cellular circuits and activities. Compared to chemical inducers, light-dependent control of gene expression by optogenetics achieves a higher spatial and temporal resolution. This could also prove decisive beyond basic research for manufacturing difficult-to-express proteins in pharmaceutical bioproduction. However, current optogenetic gene-expression systems limit this application in mammalian cells as expression levels and fold induction upon light stimulation are not sufficient. To overcome this limitation, we designed a photoswitch by fusing the blue light-activated light-oxygen-voltage receptor EL222 from *Erythrobacter litoralis* to the three tandem transcriptional activator domains VP64, p65, and Rta. The resultant photoswitch, dubbed DEL-VPR, allows an up to 400-fold induction of target gene expression by blue light, achieving expression levels that surpass those for strong constitutive promoters. Here, we utilized DEL-VPR to enable light-induced expression of complex monoclonal and bispecific antibodies with reduced byproduct expression, increasing the yield of functional protein complexes. Our approach offers temporally controlled yet strong gene expression and applies to both academic and industrial settings.

## Introduction

Biopharmaceuticals, such as monoclonal (mAb) and bispecific (bsAb) antibodies, are increasingly in demand as therapies for difficult-to-treat afflictions, such as cancer, by promoting e.g., the specific recruitment of T or NK cells to cancerous cells (1). While biopharmaceuticals offer many advantages, such as the reduction of off-target and side effects, challenges remain. These include, first and foremost, high production costs, not least caused by low yields, impurities, and limited scope for automation.

In contrast to classic IgG antibodies, bsAb do not possess two identical antigen binding-sites but two different ones. However, the production of bsAb with two different light (LC) and heavy (HC) chains leads to high levels of undesired by-products (e.g., 2 LC + 2 HC with the same antigen binding site) (2, 3). Hence, single-chain variable fragments (HCscFv) are often used, containing the variable fragments of the LC and HC. Together with other technologies, such as knobs-into-hole mutations (45), this reduces mispairing in bsAb production, but they do not completely prevent it. Static or temporal imbalances in the amounts of produced single chains contribute to the high costs of bsAb production.

One avenue towards overcoming these restrictions is the precise timing and modulation of gene expression to improve protein production, quality, and yield. Chemical induction systems, such as tetracycline- or cumate-based systems, are commonly used in research and production (5–7). However, these systems are also costly, affect cell viability, and lack temporal and spatial resolution.

The current issues in gene induction can be addressed by optogenetic induction of gene expression. Optogenetics enables the control of diverse cellular signaling processes, e.g., membrane potential, cellular signaling pathways, and gene expression by light (8–11). Systems regulated by blue light show high versatility, efficiency and favorable kinetics (12). Notably, blue-light-responsive photoswitches have been advanced for regulating gene expression in a variety of organisms, spanning bacteria (13–16), yeast (17), and mammals (18, 19). Setups for the optogenetic induction of gene expression generally rely on light-activated protein-protein and protein-DNA interactions (12, 20). In mammalian cells, blue light-activated optogenetic systems are mostly based on either plant cryptochromes (21, 22) or light-oxygen-voltage domains (LOV) (23–25). Although receptors of the phytochrome superfamily have also been deployed for optogenetic gene expression control in eukaryotic cells (26), the pertinent systems suffer from requiring reduced bilin chromophores that are not available in mammals. In contrast, both cryptochromes and LOV receptors resort to flavin-nucleotide chromophores, which are ubiquitously abundant as redox-active cofactors. Compared to cryptochrome and phytochrome-based systems, which generally need at least two different components for photo-sensing, certain LOV receptors rely on a single component and act by forming a light-induced homodimer (26–28). This makes the system less prone to variations in transfection efficiency and cellular expression levels. Moreover, it reduces the genetic footprint of the optogenetic circuit and saves valuable space in expression vectors, especially when using viral vectors.

The LOV protein EL222, deriving from *Erythrobacter litoralis* HTCC2594 (29), consists of two functional domains, the light-sensitive LOV domain and a helix-turn-helix (HTH) DNA-binding domain. In the resting state, the HTH 4α helix is covered by the LOV domain, which in turn prevents dimerization and DNA binding (29). Blue light induces the formation of a metastable covalent adduct between a cysteine residue of the LOV domain and flavin mononucleotide (FMN), which serves as the light-sensing cofactor (24, 30, 31). The concomitant protonation of the flavin N5 atom triggers conformational changes (32) and results in the release of the HTH domain, which leads to receptor dimerization and DNA binding of the HTH domain. (33, 34). To achieve an optimized DNA binding of the EL222-binding sequence, Clone 1-20 bp (C120) with a downstream TATA box promoter is often used for gene induction (28, 33). The LOV system has activation kinetics of only a few seconds with a spontaneous reversion in the dark (ca. 50 s at ambient temperature) (28, 33). In previous studies, EL222 only showed minor basal activity of non-induced gene expression as well as limited cytotoxicity (28, 33, 35).

Modular photoswitches have been developed using EL222 as the sensing domain, of which two, VP-EL222 and VEL, are particularly attractive (28, 36). These tools represent consecutive generations of engineered transactivators, which have been adapted for high-level protein expression in mammalian cells. VP-EL222 is characterized by one nuclear localization sequence (NLS), while VEL, which features an optimized EL222 sequence, contains two NLSs. Both also incorporate the activation domain VP16 from the herpes simplex virus (HSV) 1 (37), which, however, is a comparatively weak and slow-acting activation domain (38).

To overcome the limitations of the current systems and achieve a fold induction of light-mediated gene expression that allows the expression of biotherapeutics like antibodies in a sufficient yield and quality, we combined the EL222 constructs with the transactivating domains VPR (39–41), and thereby engineered the new blue-sensitive modular photoswitch DEL-VPR. VPR is a fusion product of VP64 (four repeats of the VP16 core) and the transactivation domains of the NF-kappa B subunit p65 (42) and the Epstein-Barr virus R transactivator (Rta) (43). DEL-VPR contains the N-terminal SV40 NLS and the C-terminal nucleoplasmin NLS.

In this study, we show that DEL-VPR supports a high-fold induction of protein expression in HEK293T and CHO-K1 cells under blue light, reaching the level of CMV promoter-driven constitutive protein expression combined with a low basal activity. These favorable traits allowed the expression of biotherapeutics such as monoclonal and bispecific antibodies with high yield and purity.

## Material & Methods

### Plasmids

A gene encoding VP-EL222 was synthesized with mammalian codon usage by Integrated DNA Technologie (IDT), while the VEL construct and VPR sequence were synthesized by GeneArt (ThermoFisher). Later, VP-EL222 and VEL were singularly cloned inside a pcDNA3.1 vector (Invitrogen) using *BamHI* and *XbaI* restriction sites. For the light-induced expression of firefly luciferase, the constructs pGL4.23_5xC120-minP-FLuc and pcDNA3.1_5xC120-minP-FLuc were generously provided by Kevin Gardner (CCNY, USA). The construct 5xC120-minP-mCherry, used for light-induced expression of the fluorescent reporter mCherry, was created by cloning PCR-amplified mCherry into pcDNA3.1_5xC120-minP-FLuc vector using *EcoRI* and *XbaI* restriction sites. Primers are described in the Supplementary Table 1.

As a model for monoclonal IgG antibody (mAb) expression, we chose the tetrameric Trastuzumab/Herceptin, consisting of two light (LC) and two heavy (HC) chains, which was generously provided by Andrew Beavil (Addgene #61883). For the light-induced expression, the mAb chains were amplified via PCR and subcloned into pcDNA3.1_5xC120-minP-FLuc vector using *AscI* and *MreI* restriction sites. For the constitutive expression, the mAb chains were PCR-amplified and subcloned into pcDNA3.1 vector under the control of the CMV promoter, utilizing either *AscI* and *XhoI* or *AscI* and *MreI* restriction sites.

The plasmid encoding the bsAb was generously provided by Lonza. The expressed protein is a trimeric IgG bsAb with an LC and HC on one side and a single chain variable chain (HCscFv) on the other (proprietary to Lonza). The formation of homodimers of the HC or the HCscFv is limited by the knobs-into-holes technology. For constitutive expression, all three bsAb chains were encoded on the same plasmid, individually driven by a CMV promoter. For the light-induced expression of the different bsAb chains, the single chain DNA sequences (LC, HC, and HCscFv) were synthesized by GeneArt (ThermoFisher) and inserted into the pcDNA3.1_5xC120-minP vector ordered at Genscript. For the constitutive expression of the HCscFv, the synthesized sequence was inserted inside a pInducer20 vector under the control of the H1 promoter.

### Cell lines

HEK293T and CHO-K1 were obtained and authenticated from the American Type Culture Collection (ATCC).

HEK293T cells were maintained in DMEM (Gibco, #10564011) and 10% FCS (Biochrome) at 37 °C and 5% CO2. CHO-K1 cells were maintained in DMEM/F12 1:1 (Gibco, #11320033) with 10% FCS at 37 °C and 5% CO2.

### Transfection

For both the luciferase assays and the live-cell monitoring, 30, 000 HEK293T cells/well were seeded in 50 µl DMEM+10% FCS/well in two 96-well plates. One well was exposed to blue light, whereas the other well was kept in the dark. 1 h after seeding, cells were transfected with 0.1 µg DNA/well using Lipofectamine 3000 (Invitrogen, #L3000001) in a ratio of 1:3 (DNA:Lipofectamine). For the luciferase assays, the following genes were transfected in the ratio 5:1:0, 1: blue light-sensitive photoswitches. (VP-EL222, VEL, or DEL-VPR), a light-sensitive luminescent reporter ((C120)5-firefly luciferase), and constitutive luminescent reporters (SV40-renilla luciferase for the normalization of the firefly measures and HSV TK/SV40/CMV/H1-firefly luciferase as references for constitutive protein expression). For live-cell monitoring, the following genes were transfected in the ratio 5:1: blue-sensitive photoswitches (VP-EL222, VEL, or DEL-VPR), a light-sensitive fluorescent reporter ((C120)5-mCherry), constitutive luminescent reporters (CMV/H1-mCherry) as references for constitutive protein expression. Cells were transfected 8 h before light induction.

For antibody expression, cells were seeded at 140, 000 cells/well in a 24-well plate (Sarstedt) 5 h before transfection in OptiMEM (Gibco, #11058021). The HEK293T were transfected with 500 ng DNA using PEI (Sigma #408727) in a DNA to PEI ratio of 1:2.CHO-K1 cells were transfected with 500 ng total DNA using Lipofectamine3000. For mAb experiments the DEL-VPR, C120-LC, and C120-HC-encoding plasmids were transfected in a ratio of 3:1:1. For the bsAb experiments, the DEL-VPR, C120-LC, C120-HC, and C120-HCscFv-encoding plasmids were transfected in a ratio of 3:1:1:1. The cells were transfected 24 h before light induction.

### Optogenetic stimulation

Cells were illuminated at 470 nm (always ON) at the indicated intensities (µW/cm^2^) for the indicated duration using the MLA-1 multiformat illumination device (Ningaloo Biosystems) at 37°C and 5% CO2. The dark controls were always kept in the dark.

### Luciferase Assay

Cells were excited with constant blue light (470 nm) for 9 h, starting 8 h after transfection, and a light intensity of 1500 µW/cm^2^. After illumination, the dual luciferase assay was performed using the Dual-Glo® kit (Promega, Cat. Nr. E2940), according to the manufacturer’s protocol. Briefly, the culture medium was removed and substituted with 40 µL/well Opti-MEM, followed by the addition of an equal volume of Dual-Glo® Reagent and incubation for 30 minutes to promote cell lysis. Afterwards, the firefly luminescence was measured with a FLUOstar Omega microplate reader (BMG). Next, the firefly luminescence was quenched by adding one volume of Dual-Glo® Stop & Glo® Reagent, followed by 30 min incubation, after which the renilla luminescence was measured. The ratio between the firefly and the renilla luminescence allowed to normalize the measured values.

### Live-cell imaging

The light-exposed samples were excited with constant blue light (470 nm) initially for 8 h, starting 8 h after transfection, and a relative light intensity of 1500 µW/cm^2^. Then the samples were moved back and forth every 4 h for a period of 8 h between the blue light source and the IncuCyte reader (Sartorius) to allow the acquisition of the phase and the red channels. After 8 h, the samples were illuminated for an additional 16 h without any image collection. Ultimately, the samples were placed back into the IncuCyte reader for 24 h, where pictures were collected again at intervals of 4 h. The dark samples were kept inside the IncuCyte in the absence of light for the entire experiment. The images collected with the IncuCyte reader were analyzed using the IncuCyte analyzer software (IncuCyte^®^ Base Analysis Software). Consistent image processing parameters were applied across all the experiments to both the phase channel, for measuring cell confluency, and to the red channel, for quantifying the number of fluorescent cells per mm^2^ and their integrated intensity (OCU x µm^2^ / mm^2^).

### Western blot

Cell culture supernatants were collected 24 h after light induction and cellular debris was removed. A non-reducing loading dye (200 mM Tris/HCl pH 6.8, 8% (w/v) SDS, 50% (v/v) glycerin, 0.04% bromophenol blue) was added to the samples for non-reducing Western blots, which additionally contained 4% (v/v) β-mercaptoethanol for reducing Western blots. Samples were denatured for 5 min at 95 °C and loaded on precasted 4-20% SDS-PAGE (Biotrend, #DG101-02-V2). Proteins were blotted on PVDF membranes (Merck Millipore, #T381.1) using a semi-dry blotter (BioRad, #T788.1). Membranes were blocked in 5% BSA (Sigma-Aldrich # A2153). Produced antibodies were detected with a goat-anti-human IgG antibody coupled to HRP (Thermo Fisher; 1:5000) and ECL (Perkin Elmer, #10400505). The band intensities were quantified using Fiji (44).

### ELISA

Crude cell culture supernatants were collected 24 h after light induction. Antibody production was detected using the ELISA human IgG total kit (Thermo Fisher, #88-50550-22) according to the manufacturer’s protocol. The absorption was measured using a plate reader (Tecan Infinite 200 pro). Antibody expression was calculated in pg/cells/day.

### Statistical analysis

Data were plotted using Microsoft 365 Excel. Microsoft Excel and JASP (JASP Team (2024), version 0.19.0 [Windows 64 bit]) were used to perform statistical analysis. Before determining the P values between independent samples, the normality of their distribution was verified with the Shapiro-Wilk test. If passed, the Student’s t-test was performed and if not, the Mann-Whitney test. The one-way ANOVA followed by the Holm-Sidak *post-hoc* test was performed for comparing more than two groups with one independent variable, and for more than two groups with two independent variables the two-way ANOVA with Holm-Sidak post-hoc test. The data is represented as mean ± SD of n = 5 (Western Blot, ELISA) or n = 4 (luciferase assay, live-cell imaging) experiments. The single dots display the values of individual samples (Western Blot) or of triplicates.

## Results

### Design of the modular blue-light sensitive photoswitch DEL-VPR

To improve gene expression control for protein expression, e.g., antibody production, we aimed to develop an optogenetic tool for precise temporal control over gene activation and concomitantly enhanced expression levels, while promoting functional protein complex formation (Figure 1A). We identified the bacterial EL222-based expression systems as promising candidates due to their compact size, natural occurrence of the chromophore, and proven performance. We tested the already published EL222-based systems (28) using a luciferase reporter assay, and demonstrated that while functional, the overall expression levels were rather low (Figure 1B). Thus, we aimed to improve these photoswitches by achieving higher expression levels during the turn-on phase, while minimizing leakiness during the turn-off phase, based on the expression of a single component with minimal basal activity in the dark. To this end, we engineered a new blue-light sensitive transactivator, DEL-VPR, based on the bacterial sensor EL222 (28), combined with three distinct activation domains (VP64, p65, and Rta), connected by flexible GS linkers (Figure 1A).

**Figure 1:**
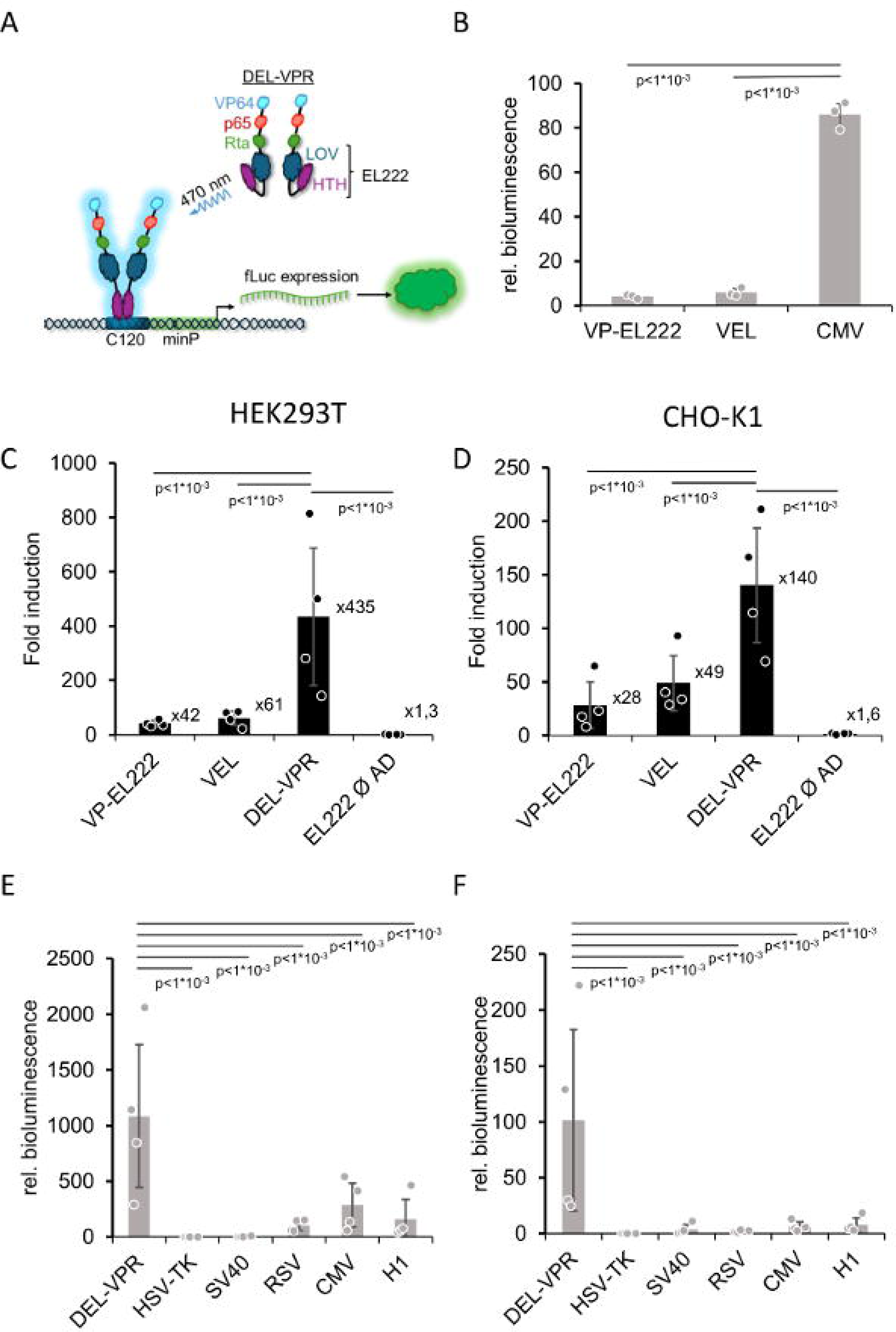
Light-dependent luciferase activity in HEK293T and CHO-K1 cells. **A)** DEL-VPR is activated by blue light (470 nm) and dimerizes and binds to the C120 sequence. Subsequently, firefly luciferase (fLuc) is expressed in a light-dependent manner. B) Initial experiments were performed in HEK293T cells, comparing the performance of published EL222-based photoswitches with the constitutive CMV promoter. C-D) Luminescence levels were measured in HEK293T (C) and CHO-K1 (D) cells, which transiently expressed fLuc either in a light-dependent fashion or under different constitutive promoters. Different versions of the blue-sensitive photoswitch EL222 were compared (VP-EL222, VEL, DEL-VPR, and EL222 without additional activation domain) after being excited for 9 h using blue light (470 nm) with a constant light intensity (1500 µW/cm^2^). Data represent the fold change of the means of firefly luminescence values detected in samples exposed to light versus dark, normalized by their corresponding renilla luminescence values. E-F) Comparison of samples where expression was either induced by light through the activation of the transactivator DEL-VPR or maintained continuously under the control of one of the following promoters (HSV-TK, SV40, RSV, CMV, or H1). Data are presented as bar graphs, showing mean SD, and dots represent individual samples. Statistical analysis (D-E) was performed using a one-way ANOVA with the Holm-Sidak post-hoc test. In B), plots contain one experiment with three samples, while in E-F) four (n = 4) independent experiments, each representing the means of three samples (total n =12).

To assess the efficiency of DEL-VPR, we compared it to alternative versions of EL222 and common constitutive promoters using a luciferase reporter assay (Figure 1—Supplementary Figure 1). To this end, we transfected HEK293T or CHO-K1 cells with either an EL222-based blue light-sensitive system (VP-EL222, VEL, EL222 without additional activation domain, or DEL-VPR) and the light-inducible reporter-gene cassette, or only with a plasmid driving the reporter expression via a constitutive promoter (HSV-TK, SV40, RSV, CMV, or H1). DEL-VPR dramatically outperformed both the VP-EL222 and VEL transactivators, resulting in a factor of more than 400-fold between expression levels for blue light and dark in HEK293T (Figure 1C—Supplementary Figure 1A) and 140-fold in CHO-K1 cells (Figure 1D— Supplementary Figure 1B). Furthermore, the light-induced luciferase expression driven by DEL-VPR was significantly higher compared to the expression levels mediated by weak and medium constitutive promoters like HSV-TK and SV40, with differences ranging from 2500- to 1000-fold in HEK293T cells (Figure 1E— Supplementary Figure 1C) and 1200- to 60-fold in CHO-K1 cells (Figure 1F— Supplementary Figure 1D). The expression levels achieved by DEL-VPR upon light excitation were also comparable to those driven by strong constitutive promoters like RSV, CMV, and H1 with differences of 10- to 5-fold in HEK293T cells (Figure 1E— Supplementary Figure 1C) and 50-to 15-fold in CHO-K1 and CHO-K1 cells (Figure 1F—Supplementary Figure 1D). Together, these data not only underline the superior performance of the DEL-VPR transactivator in comparison to previously published EL222 variants but also show that the maximal light-induced expression levels can compete with those of very strong constitutive promoters.

### Time course of light-induced protein expression

Next, we investigated the performance of DEL-VPR in driving the expression of the fluorescent reporter mCherry in live cells (Figure 2—Supplementary Figure 2). We compared its efficiency to the alternative EL222 versions VP-EL222 and VEL (Figure 2C-D—Supplementary Figure 2B-C) and the strong constitutive CMV promoter (Figure 2E-F—Supplementary Figure 2D-E). After light activation, DEL-VPR effectively mediates mCherry expression at levels comparable to those induced by the CMV promoter, without any detectable basal expression in the dark (Figure 2A, B—Figure 2C-D bottom panels—Supplementary Figure 2B-C bottom panels). mCherry fluorescence increased during light stimulation (up to the 24 h time point, Figure 2—Supplementary Figure 2, marked with “+”) and then remained stable post-stimulation. In HEK293T cells, the proportion of mCherry-positive cells induced by DEL-VPR was similar to that achieved by the CMV promoter (Figure 2E), although the average fluorescence intensity was significantly higher in CMV-driven cells (Supplementary Figure 2B, D). In CHO-K1 cells, mCherry expression was lower and more variable compared to HEK293T cells, though similar trends were observed like DEL-VPR inducing expression levels comparable to those of the CMV promoter (Figure 2D, F—Supplementary Figure 2C, E). Notably, VEL also showed comparable mCherry-positive cell counts and fluorescence intensity to DEL-VPR (Figure 2D— Supplementary Figure 2C). Still, both systems significantly enhanced mCherry expression following light activation relative to the dark (Figure 2B, D bottom panel— Supplementary Figure 2C bottom panel). Despite its relatively lower performance in CHO-K1 cells, DEL-VPR remains an efficient alternative to canonical constitutive expression systems, especially when considering its dynamic light-controlled modulation and the absence of detectable basal mCherry expression in the dark.

**Figure 2:**
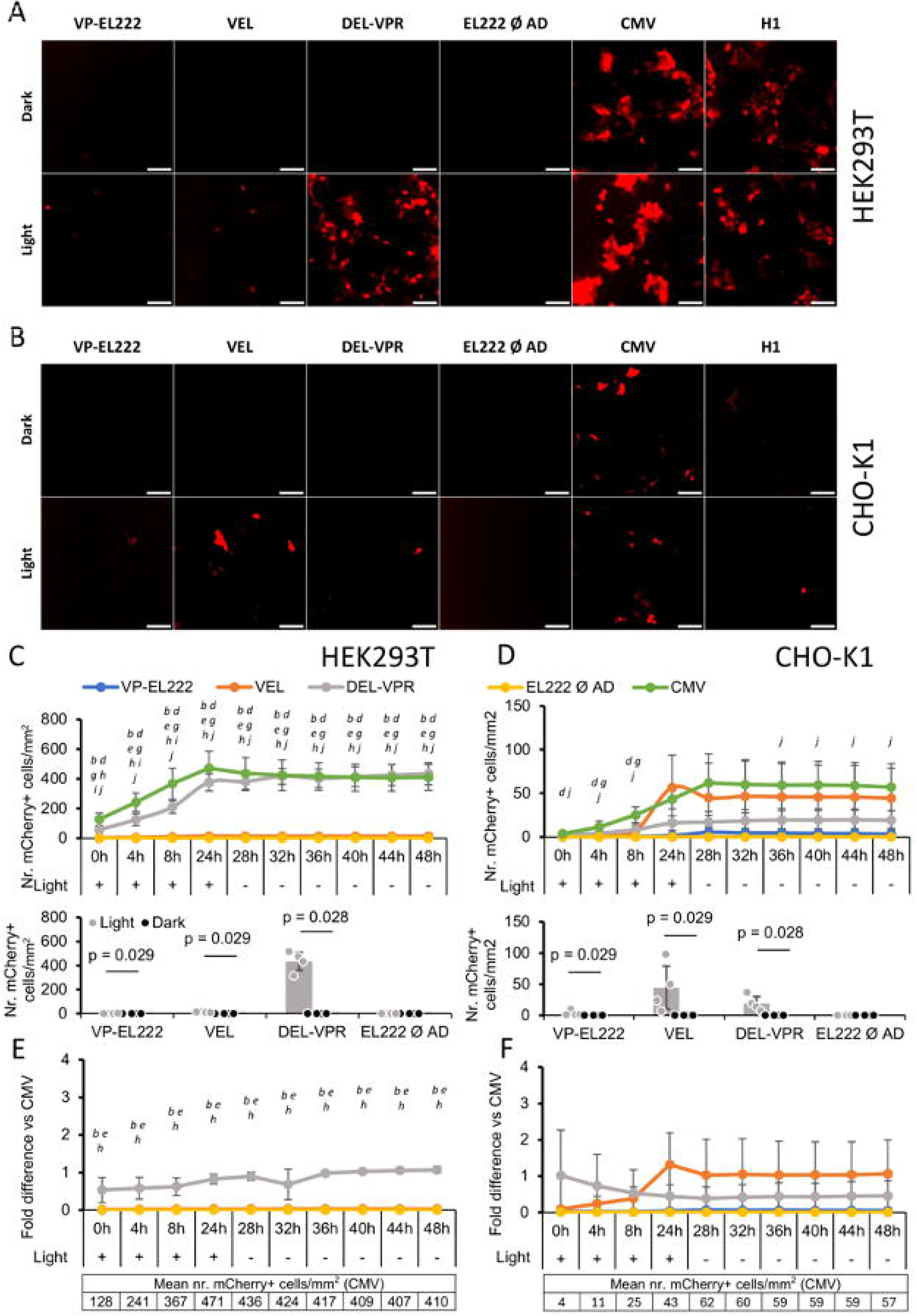
Light-dependent fluorescent reporter expression in HEK293T and CHO-K1. Fluorescence levels were measured in HEK293T (A, C, E) and CHO-K1 (B, D, F) cells, which transiently expressed mCherry either in a light-dependent fashion or under the control of different constitutive promoters. We compared different versions of the blue-sensitive photoswitch EL222—VP-EL222, VEL, DEL-VPR and EL222 without additional activation domain—and the strong and medium constitutive promoters CMV and H1, respectively. For the light condition, samples were excited for 32 h using blue light (λ= 470 nm) with a constant intensity (1500 µW/cm2), while for the dark condition, samples were kept in the dark for the entire time. Live cell image collection started 8 h after the beginning of the light excitation (+) and continued for a total of 48 h. For the last 24 h, all the samples were kept in the dark (-). In A) and B) are shown examples of the images collected and analyzed. C) and D) top panels represent the time course of the mean number of mCherry fluorescence positive cells per mm2 detected in samples exposed to the light, while the bottom panels show the mean values relative to the last time point (48 h) of the light and the dark conditions compared. The graphs E) and F) represent the fold difference of the mean number of mCherry fluorescent cells transfected with the different versions of the photoswitch EL222 versus the one detected in the samples transfected with the strong constitutive promoter CMV. Data are presented as lines with markers or as bar graphs, showing mean ± SD, and with dots indicating individual values of each sample. Statistical analysis was performed in C), D), E) and F) using a one-way ANOVA with Holm-Sidak post-hoc test. Statistically significant differences among the different photoswitches with p<0, 05 were indicated as follows: *a*=VP-EL222 vs. VEL; *b*=VP-EL222 vs. DEL-VPR; *c*=VP-EL222 vs. EL222 Ø AD; *d*=VP-EL222 vs. CMV; *e*=VEL vs. DEL-VPR; *f*=VEL vs. EL222 Ø AD; *g*=VEL vs. CMV; *h*=DEL-VPR vs. EL222 Ø AD; *i*=DEL-VPR vs CMV; *j*=EL222 Ø AD vs CMV. In the bottom panels of C) and D), the statistical analysis was performed using the Mann-Whitney test after having verified the distribution with a Shapiro-Wilk test. Plots contain four (n=4) independent experiments, each representing the means of three samples (total n=12).

### Optogenetic production of a monoclonal antibody

As our data pinpointed DEL-VPR as a powerful tool for modulating gene expression, we investigated whether more complex proteins, such as monoclonal antibodies (mAb), widely used in biotherapeutics, could also be efficiently produced using this optogenetic strategy. We chose the human epidermal growth factor receptor 2 (HER2)-targeted mAb Trastuzumab (IgG), also known as Herceptin, which is used to treat HER2-positive breast and gastric cancer by inhibiting the HER2 signaling pathway and thereby limiting proliferation (45, 46). The mAb comprises two light (LC) and two heavy (HC) chains that assemble into a heterotetramer. We cloned the LC and HC downstream of the light-regulated promoter C120 and co-expressed the two chains under the control of DEL-VPR (Figure 3A). In both HEK293T and CHO-K1 cells, mAb expression was not detectable in the dark (Figure 3B-E). However, blue-light excitation increased mAb expression, assembly, and secretion into the supernatant in a light dose-dependent manner (Figure 3B-E). In HEK293T cells, mAb expression was maximally induced at 1500 µW/cm^2^ blue light (BL), while in CHO-K1 cells, it was already reached at 1000 µW/cm^2^ BL and slightly dropped again at 1500 µW/cm^2^ BL. At the maximal blue-light intensity, the expression levels reached or exceeded those afforded by the constitutive CMV promoter.

**Figure 3:**
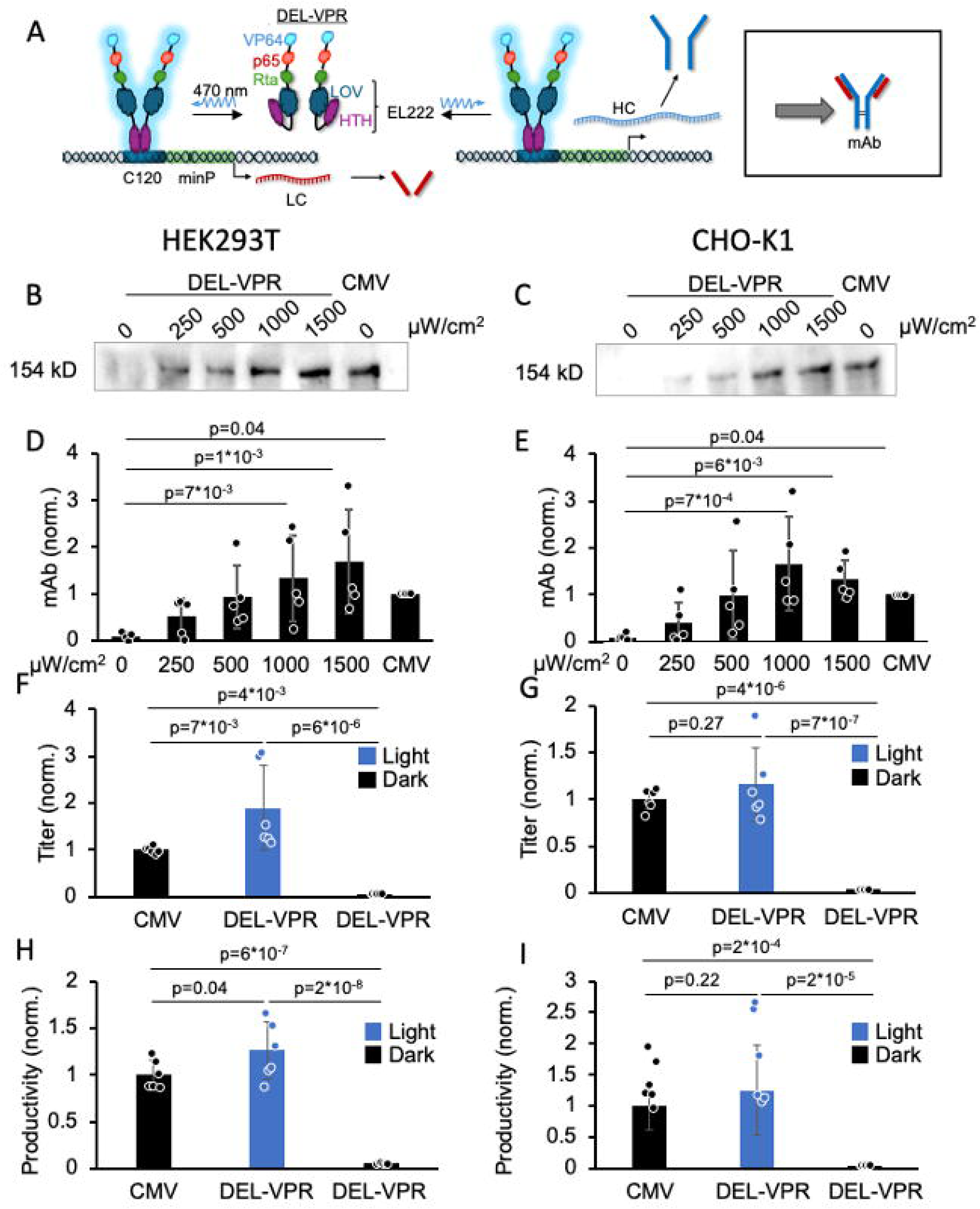
Light-dependent titration of mAb expression in HEK293T and CHO-K1. A) DEL-VPR is activated by blue light excitation (λ = 470 nm) and in turn, dimerizes and binds to the C120 sequence. Subsequently, the light (LC) and heavy (HC) chains are expressed in a light dose-dependent manner, followed by their assembly into mAb in the ER. The mAb, consisting of the light and heavy chains, was either expressed in HEK293T (B) or CHO-K1 (C) either light-dependently under the control of the C120 promoter or constitutively under the control of the CMV promoter. The light-dependent conditions were excited at different intensities (µW/cm^2^) for 24 h (always ON). The samples were analyzed on a Western blot (WB) under non-reducing conditions using an anti-human IgG1 detection antibody coupled to HRP. The mAb bands of the WB for the HEK293T and CHO-K1 were quantified in D) and E), respectively. F) and G) show the fold change of mAb titers of the light-induced (DEL-VPR light; 1500 µW/cm^2^), non-induced (DEL-VPR dark) or the constitutive (CMV) conditions after 24 h expression in HEK293T or CHO-K1, respectively, quantified by ELISA. The values were normalized to the CMV values. H) and I) show the fold change of mAb productivity of the experiment presented in F-G) in HEK293T and CHO-K1, respectively. The values were normalized to the CMV values. D-I) Means of biological independent samples ± STD are presented as bars, dots indicate individual values of each sample. Statistical analysis was performed using one-way ANOVA with the Holm-Sidak post-hoc test. Plots contain five (n=5) in D-E) and six (n=6) independent samples in F-I).

To determine the antibody titer, we performed an ELISA against human IgG demonstrating a significant increase upon light induction by DEL-VPR in HEK293T and CHO-K1, while in the dark the titer barely exceeded the value of the negative control (Figure 3F-G—Supplementary Figure 3A, B). The same holds true for the productivity measured in pg/cell/day (Figure 3H, I—Supplementary Figure 3C-D). The cells were incubated in the dark for the first 24 h after transfection, only allowing the expression of proteins under a constitutive promoter (CMV-DEL-VPR, CMV-mAb). Then, the light-dependent proteins were produced for 24 hours. Accordingly, the constitutive proteins were produced for a total of 48 h, whereas the DEL-VPR-induced proteins were only produced for 24 h. Even despite the shorter production time, the mAb titer and productivity were significantly increased by DEL-VPR compared to the constitutive CMV-driven expression in HEK293T (Figure 3F, H— Supplementary Figure 3A, C). However, in CHO-K1 cells, the titer and productivity were not significantly different between the DEL-VPR (light) and CMV (Figure 3G, I—Supplementary Figure 3B, D). Thus, our data demonstrate that DEL-VPR allows a tightly controlled, light-dependent induction of mAb expression, reaching or even surpassing the level of strong constitutive expression mediated by a CMV promoter.

### Optogenetic production of a bispecific antibody

Having successfully demonstrated the light-dependent expression of mAb, we next applied our approach to more complex biopharmaceuticals like a bispecific antibody (bsAb). Since one of the driving factors for the high production costs of bsAb is a contamination of unwanted by-products (e.g. HCscFv homodimers), we investigated whether we could modulate bsAb complex assembly by light-mediated induction. To establish this, we initially tested the light regulation of all bsAb chains simultaneously.

We chose a trimeric bsAb (IgG), comprising an LC and HC on one side and an HCscFv on the other side. We cloned the three constituent bsAb chains downstream of the light-responsive promoters C120 and co-expressed them with DEL-VPR (Figure 4A).

**Figure 4:**
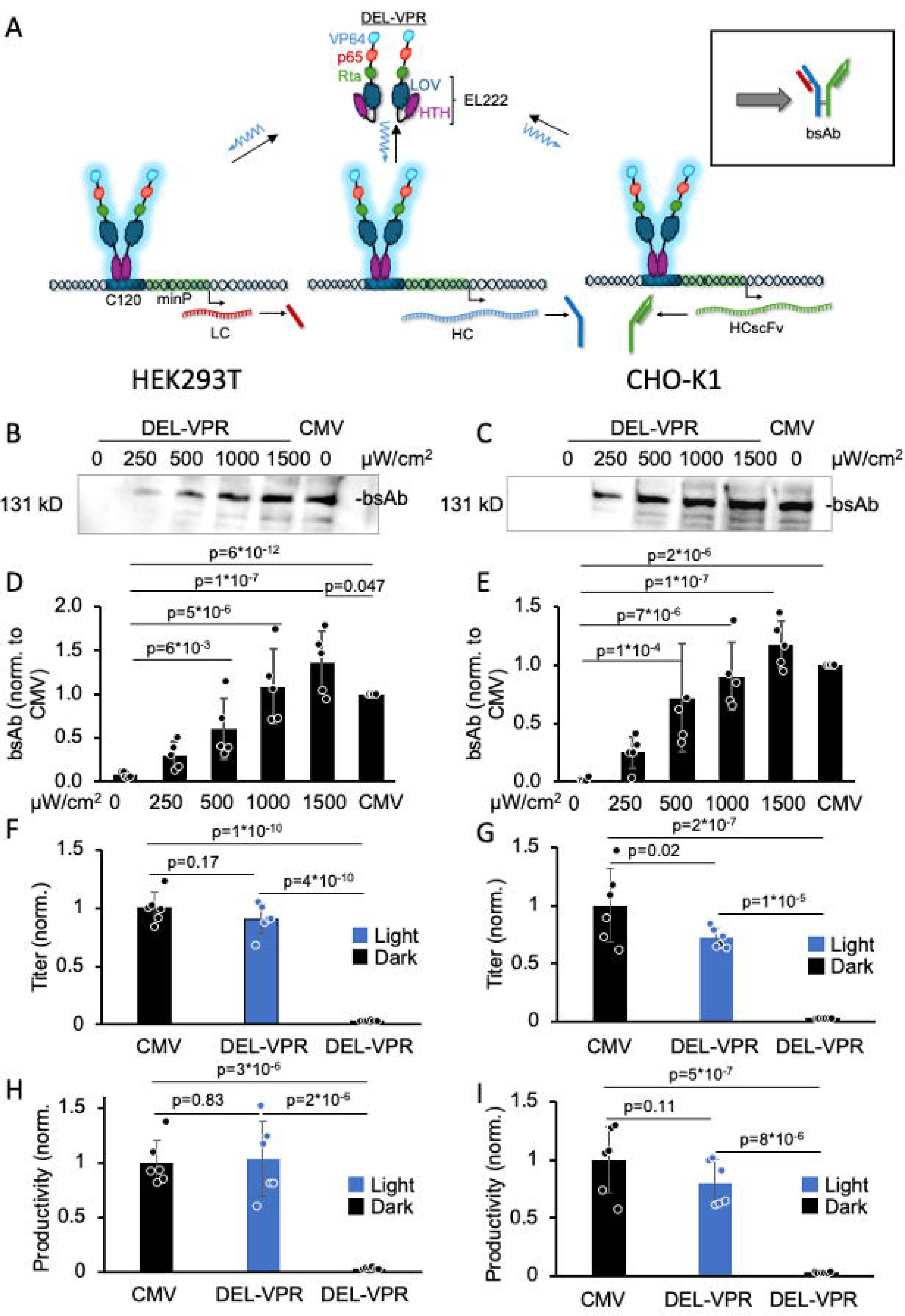
Light-dependent titration of bsAb expression in HEK293T and CHO-K1. A) DEL-VPR is activated by blue light excitation ( λ= 470 nm) and in turn, dimerizes and binds to the C120 sequence. Subsequently, the light (LC), heavy (HC) and single variable fragment (HCscFv) chains are expressed in a light dose-dependent manner, followed by their assembly into bsAb in the ER. The bsAb was either expressed in HEK293T (B) or CHO-K1 (C) either light-dependently under the control of DEL-VPR or constitutively under the control of the CMV promoter. Cells were excited at different intensities (µW/cm^2^) for 24 h (always ON). The samples were analyzed on a Western blot (WB) under non-reducing conditions using an anti-human IgG1 detection antibody coupled to HRP. The bsAb bands of the WB for the HEK293T and CHO-K1 were quantified in D) and E), respectively. F) and G) show the fold change of bsAb titer of the light-induced (DEL-VPR; 1500 µW/cm^2^), non-induced (DEL-VPR dark) or constitutive (CMV) conditions after 24 h expression in HEK293T or CHO-K1, respectively, quantified by ELISA. The values were normalized to the CMV samples. H) and I) show the fold change of bsAb productivity of the experiment presented in F-G) in HEK293T and CHO-K1, respectively. The values were normalized to the CMV samples. D-I) Means of biological independent samples ± STD are presented as bars, dots indicate individual values of each sample. Statistical analysis was performed using one-way ANOVA with the Holm-Sidak post-hoc test. Plots contain five (n=5) in C-E) and six (n=6) independent samples in F-I).

In both HEK293T and CHO-K1 cells, bsAb expression using DEL-VPR and secretion was barely detectable in the dark but significantly increased in a light dose-dependent manner, as observed by trimeric bsAb complex assembly in the supernatant (Figure 4B-E), for both HEK293T and CHO-K1 cells, maximal bsAb expression was observed at 1500 µW/cm². In HEK293T cells, the bsAb level was significantly higher in the DEL-VPR samples (1500 µW/cm² BL) compared to the CMV-bsAb control (Figure 4B, D). In CHO-K1 cells, the DEL-VPR-induced samples showed a tendency towards higher expression than the CMV samples (Figure 4C, E). Strikingly, DEL-VPR-mediated expression resulted in the predominant assembly of the trimeric bsAb in the supernatant, whereas the other conditions displayed more undesired byproducts.

DEL-VPR led to a significant induction of the bsAb titer under BL compared to the dark by 30-fold and 34-fold while increasing the productivity by 32-fold and 33-fold in HEK293T and CHO-K1, respectively (Figure 4F-I—Supplementary Figure 4). The antibody titer was not significantly different between the DEL-VPR-induced and CMV-driven expression in HEK293T cells (Figure 4F), whereas in CHO-K1 cells, the bsAb levels were slightly higher in the CMV-driven expression compared to the DEL-VPR-driven expression (Figure 4G—Supplementary Figure 4B). The productivity in pg/cell/day did not differ between the constitutive and the DEL-VPR-induced expression (Figure 4H, I—Supplementary Figure 4C-D). Of note, the constitutive expression occurred over 48 h, whereas the DEL-VPR-induced expression only occurred during the last 24 h of the experiment.

In summary, these data demonstrate that DEL-VPR can be used as an optogenetic tool for the tunable expression of large protein complexes, reaching levels of constitutive protein expression driven by, e.g., a CMV promoter but reducing the formation of undesired byproducts. By allowing to switch on bsAb expression during a particular desired time window, DEL-VPR stands to reduce the metabolic burden that can occur with a static high expression using constitutive promoters such as CMV.

### bsAb complex composition can be modulated by light

When we induced all three bsAb chains at the same time (Figure 4B-E), we observed the trimeric bsAb as the predominant product. In addition to regulating the production of the bsAb as such, we examined whether the composition of the bsAb complexes could be modified by light since by-products represent a major factor impeding purification and yield in bsAb production. To this end, we constitutively expressed the HCscFv chain combined with the light-dependent expression of LC/HC driven by DEL-VPR (Figure 5A-B). This enables a light-dependent transition from the HCscFv dimer in the dark to the trimeric bsAb complex in the presence of light, thereby establishing the proof of concept for light-dependent modulation of protein complex assembly.

**Figure 5:**
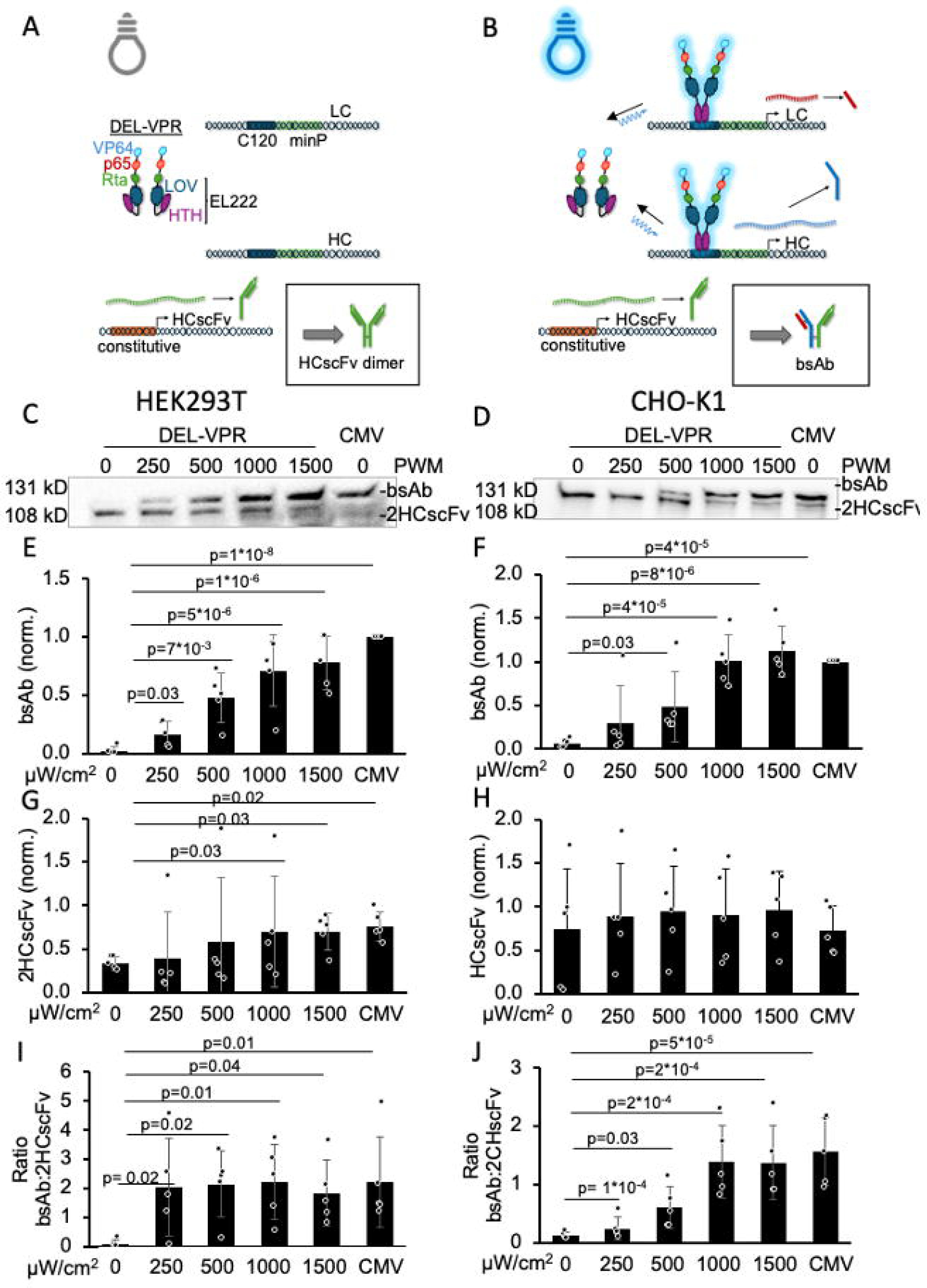
Light-dependent titration modulation of bsAb complexes in HEK293T and CHO-K1. A) In the dark, the DEL-VPR stays inactive. Therefore, the light-dependent expression of the light (LC) and heavy (HC) chain is not induced, while the single chain variable fragment (HCscFv) is constitutively expressed. Consequently, in the absence of the LC and HC, the HCscFv is assembled in dimers. B) DEL-VPR is activated by blue light excitation (λ = 470 nm) and in turn, dimerizes and binds to the C120 sequence. Subsequently, the LC and HC are expressed in a light intensity-dependent manner. Additionally, the HCscFv is expressed constitutively. Consequently, in the light, the trimeric bsAb is assembled. The bsAb was expressed in HEK293T (C) or CHO-K1 (D) either partially light-dependently (light-dependent LC/HC + constitutive HCscFv) under the control of DEL-VPR or all chains constitutively under the control of the CMV promoter. The light-dependent conditions were excited at different intensities (µW/cm^2^) for 24 h (always ON). The different bsAb complexes were analyzed on a Western blot (WB) under non-reducing conditions using an anti-human IgG1 detection antibody coupled to HRP. The bsAb bands of the WB for the HEK293T and CHO-K1 were quantified in E) and F), respectively. The bands of HCscFv dimers for the HEK293T and CHO-K1 were quantified in G) and H), respectively. The ratio of bsAb:HCscFv dimers for the HEK293T and CHO-K1 were quantified in I) and J), respectively. E-J) Means of biological independent samples ± STD are presented as bars; dots indicate the individual values of each sample. Statistical analysis was performed using ANOVA with the Holm-Sidak post-hoc test. Plots contain five (n=5) independent samples.

In the dark, only the constitutively expressed HCscFv was produced and subsequently assembled and secreted as HCscFv dimers into the supernatant in both HEK293T and CHO-K1 cells (Figure 5C-F). We observed HCscFv dimers in all DEL-VPR conditions in the dark (Figure 5C-D), but the expression of the LC and HC in a light dose-dependent manner resulted in a switch from the HCscFv dimers to the trimeric bsAb. This was also demonstrated by the ratio of bsAb to HCscFv dimer, which increased in a light dose-dependent manner (Figure 5C-D, G-H). At maximal expression (1500 µW/cm^2^ BL), both bsAb levels and the ratio of bsAb to HCscFv dimers reached levels like a CMV-driven expression (Figure 5C-F, I-J).

In conclusion, our data show that DEL-VPR allows not only dynamic regulation of the amount of biopharmaceuticals produced but also their correct assembly into protein complexes. The use of DEL-VPR, therefore, offers the possibility to dynamically modify the composition of bsAb.

## Discussion

In this study, we engineered the light-activated transcription factor DEL-VPR that allows the stringent induction of target gene expression, including monoclonal (mAb) and bispecific (bsAb) antibodies, in a light dose-dependent manner. The system is capable of induction of up to 400-fold, thereby reaching or even exceeding the levels of very strong, constitutive promoters such as CMV. Evidently, the strength of the overall expression does not have to be sacrificed at the expense of regulation by light. As DEL-VPR exhibits minimal basal activity in the dark, it could apply to the expression of toxic proteins with minimized impact on host cell viability. Thereby, DEL-VPR opens new opportunities for bioproduction, tuning the production of biotherapeutics to the desired concentration, which reduces manufacturing costs, thereby increasing the availability of these valuable therapies.

### Optogenetic tools to control gene expression

Identifying a suitable optogenetic tool to control gene expression can be challenging, although impressive progress has been made over the last decade (11). Certain blue light-based tools have gained acceptance due to their favorable induction kinetics, low basal activity, comparatively simple architecture, reliance on endogenous chromophores, and high fold induction (21, 28, 33, 47). Nevertheless, due to the novelty of this approach, optimization steps are still needed, especially concerning robustness and induction strength.

To ensure the robustness for the expression of various genes, we selected EL222 as it does not rely on binding to endogenous DNA such as, e.g., CRISPR/Cas-based tools but on the artificial promoter C120, thereby possessing consistent binding properties (28, 33, 34, 48). VP-EL222 (VP16 fused to EL222) and its modified version VEL (VP16-EL222 with an additional NLS) have previously achieved 100-fold induction with low basal activity in the dark in HEK293T (28, 33). Despite the lack of published absolute values or comparisons with constitutive promoters, our relative values for induction of gene expression after light stimulation are largely consistent with the literature using these tools. We obtained an approximately 50-fold increase in luciferase activity, but the induced absolute expression level was comparatively weak and insufficient for visualizing mCherry expression. Accordingly, we replaced the VP16 activation domain for the VPR tandem, which previously demonstrated high levels of induction in various contexts (39–41). Light-stimulated DEL-VPR resulted in a 400- and 140-fold increase in luciferase activity in HEK293T and CHO-K1, respectively. Simultaneously, DEL-VPR was responsible for a strong and highly titratable induction of fluorescent proteins and monoclonal and bispecific antibodies. At the same time, the DEL-VPR, VP-EL222 and VEL all show similar low basal activities. Consequently, DEL-VPR combines optimal properties in terms of strength (high fold induction), kinetics and tightness (low dark activity), making it an ideal candidate for modulating large-scale bioproduction.

### Advantages of optogenetic over chemical induction of gene expression

Numerous tools are available that use small molecules as inducers of gene expression (11, 49, 50). Chemically induced gene expression, e.g., by the most frequently used tetracycline/doxycycline system, is highly optimized (7), providing strong induction of up to ∼100-fold (6, 51) while having a low basal activity in the dark (52). However, there are several disadvantages associated with chemical induction. For example, off-target effects such as inhibition of mitochondrial gene expression, increased glycolysis, and slowed proliferation have been demonstrated with doxycycline (at concentrations used for induction) (53, 54).

Chemical inducers also have disadvantages that may be more significant in large-scale bioproduction than in basic research. These include, above all, the limited possibilities for automating the individual and customized inducer concentration, as well as the high costs of GMP-grade inducers. In addition, the removal of the inductors is cumbersome, and it is often necessary to replace the entire medium. This is not only extremely costly in large volumes but also results in slow deactivation kinetics. Accordingly, periodic ON/OFF regulation would not be feasible using chemical inductors. Therefore, chemical induction might be less advantageous compared to using optogenetic tools in numerous aspects of basic research and large-scale bioproduction.

## Conclusions

DEL-VPR opens new opportunities in bioproduction, namely the dynamically titrable expression of biopharmaceuticals without sacrificing the high production levels that have so far only been achieved by constitutive promoters. At the same time, the DEL-VPR could be used to reduce protein toxicity and to minimize the metabolic burden of protein overexpression since bsAb expression could be kept inactive during the selection and expansion phase. Alongside absolute expression levels, the folding and post-translational maturation of the antibodies are the main bottlenecks in limiting the yield (55). Based on the fast response times and reversibility of optogenetic regulation, the prospective enhancement of real-time feedback could facilitate the production of proteins in a manner that aligns with the cells’ capacity for their processing, thereby paving the way for the next generation of bioproduction. DEL-VPR could help to express only as much protein as can be processed. DEL-VPR’s features could help to reduce the aggregation and toxicity of non-processed antibodies and thus increase their titer.

In summary, DEL-VPR has the potential to be employed in a multitude of applications within the field of bioprocessing. These include the induction of the expression at a desired point in time, the modulation of protein complex composition and the regulation of the expression of toxic proteins. Therefore, DEL-VPR could facilitate more efficient production of various biopharmaceuticals, thereby reducing their production costs and increasing their accessibility as standard therapies for targeting difficult-to-treat conditions such as cancer and autoimmune diseases.

Beyond bioproduction, DEL-VPR supports multiple use cases in basic research and cell biology, e.g., to modulate the progression of the cell cycle, as well as to induce or prevent apoptosis, thereby determining cell survival. Furthermore, DEL-VPR could be employed to modulate signaling pathways, for example, to induce differentiation of host or stem cells for cell therapeutics or to control metabolic processes.

## Acknowledgments

We thank Kevin Gardner for providing the 5xC120-minP-FLuc construct, Andrew Beavil for providing the Trastuzumab construct and Lonza for providing the bsAb construct.

## Author contribution

DW, HMS, EC and JG designed the experiments. EC, RS, FD and JG executed the experiments. EC and JG analyzed and interpreted the data. EC and JG wrote the manuscript. DW, HMS, HMH and AM critically revised the manuscript for important intellectual content.

## Data availability

The original Western Blots and live-cell imaging picture, as well as the Excel files from the analysis, are available (https://doi.org/10.5281/zenodo.13836906).

