## Supplementary Figures for "Potent photoswitch for expression of biotherapeutics in mammalian cells by light"

**Supplementary Information**

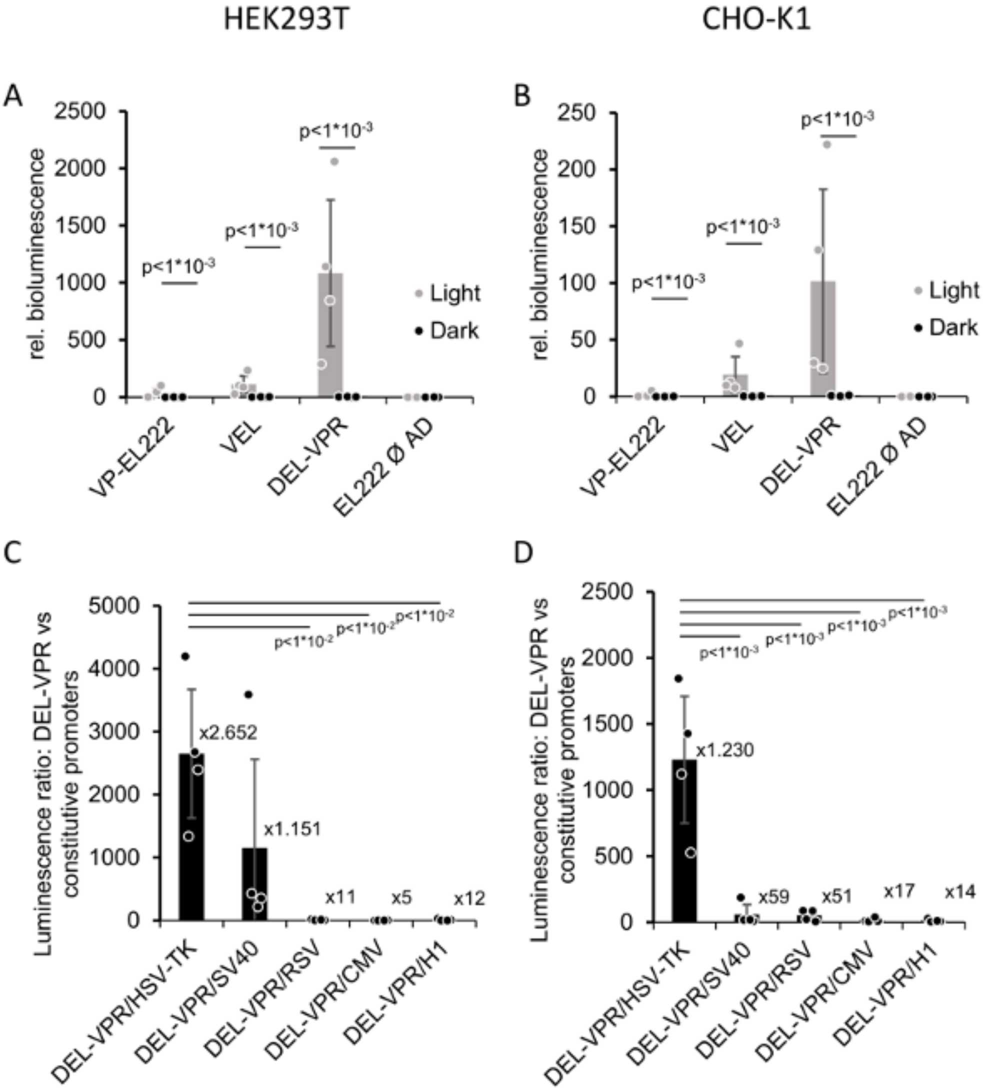


**Supplementary Figure 1: Light-dependent luciferase activity in HEK293T and CHO-K1.**

Luminescence levels were measured in HEK293T (A, C) and CHO-K1 (B, D) cells which transiently expressed firefly luciferase either in a light-dependent fashion or under the induction of different constitutive promoters. Different versions of the blue-sensitive photoswitch EL222 were compared—VP-EL222, VEL, DEL-VPR and EL222 without additional activation domain—which were excited for 9 h using blue light (λ= 470 nm) with a constant intensity (1500 μW/cm$^2$). A) and B) represent the means of firefly luminescence values detected in samples exposed to the light or kept in the dark and normalized by their corresponding renilla luminescence values. Bar graphs C) and D) illustrate the comparison of samples where expression was either induced by light through the activation of the transactivator DEL-VPR or maintained continuously under the control of one of the following constitutive promoters—HSV-TK, SV40, RSV, CMV, or H1. Means of biological independent samples $\pm$ SD are presented as bars, dots indicate the individual values of each sample. Statistical analysis was performed in A) and B) using the Mann-Whitney test after having verified the distribution with a

Shapiro-Wilk test, and in C) and D) using a one-way ANOVA with Holm-Sidak post-hoc test. Plots contain four (n=4) independent experiments, each representing the means of three samples (total n=12).

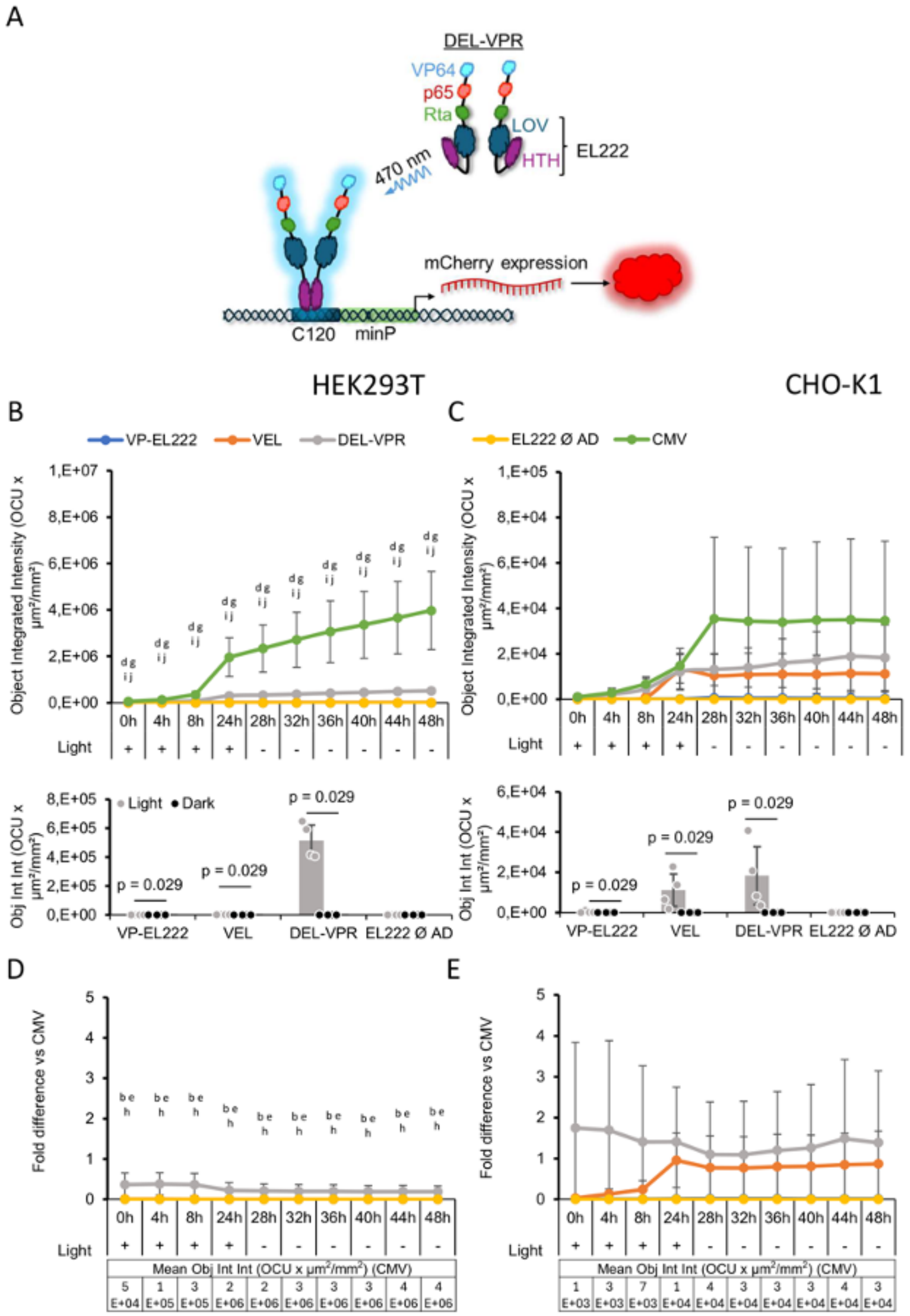


**Supplementary Figure 2: Light-dependent fluorescent reporter expression in HEK293T and CHO-K1.**

A) DEL-VPR is activated by blue light excitation (470 nm) and in turn dimerizes and binds to the C120 sequence. Subsequently, mCherry is expressed in a light-dependent manner. Fluorescence levels were measured in HEK293T (B, D) and CHO-K1 (C, E) cells which transiently expressed mCherry either in a light-dependent fashion or under the induction of different constitutive promoters. We compared different versions of the blue-sensitive photoswitch EL222—VP-EL222, VEL, DEL-VPR and EL222 without additional activation domain—and the strong and medium constitutive promoters CMV and H1, respectively. For the light condition, samples were excited for 32 h using blue light (λ= 470 nm) with a constant intensity (1500 μW/cm$^2$), while for the dark condition, samples were kept in the dark for the entire time. Live cell image collection started 8 h after the beginning of the light excitation (+) and continued for a total of 48 h. For the last 24 h, all the samples were kept in the dark (-). B) and C) top panels represent the time course of the object integrated intensity of mCherry detected in samples exposed to the light, while the bottom panels show the mean values relative to the last time point (48 h) of the light and the dark conditions compared. The graphs D) and E) represent the fold difference of the object integrated intensity of mCherry of the cells transfected with the different versions of the photoswitch EL222 versus the one detected in the samples transfected with the strong constitutive promoter CMV. Data are presented as lines with markers or as bar graphs, showing mean $\pm$ SD, and with dots indicating individual values of each sample. Statistical analysis was performed in B), C), D) and E) using a one-way ANOVA with Holm-Sidak post-hoc test. Statistically significant differences among the different photoswitches with $p<0,05$ were indicated as follow: *a*=VP-EL222 vs VEL; *b*=VP-EL222 vs DEL-VPR; *c*=VP-EL222 vs EL222 Ø AD; *d*=VP-EL222 vs CMV; *e*=VEL vs DEL-VPR; *f*=VEL vs EL222 Ø AD; *g*=VEL vs CMV; *h*=DEL-VPR vs EL222 Ø AD; *i*=DEL-VPR vs CMV; *j*=EL222 Ø AD vs CMV. In the bottom panels of B) and C), the statistical analysis was performed using the Mann-Whitney test after having verified the distribution with a Shapiro-Wilk test. Plots contain four (n=4) independent experiments, each representing the means of three samples (total n=12).

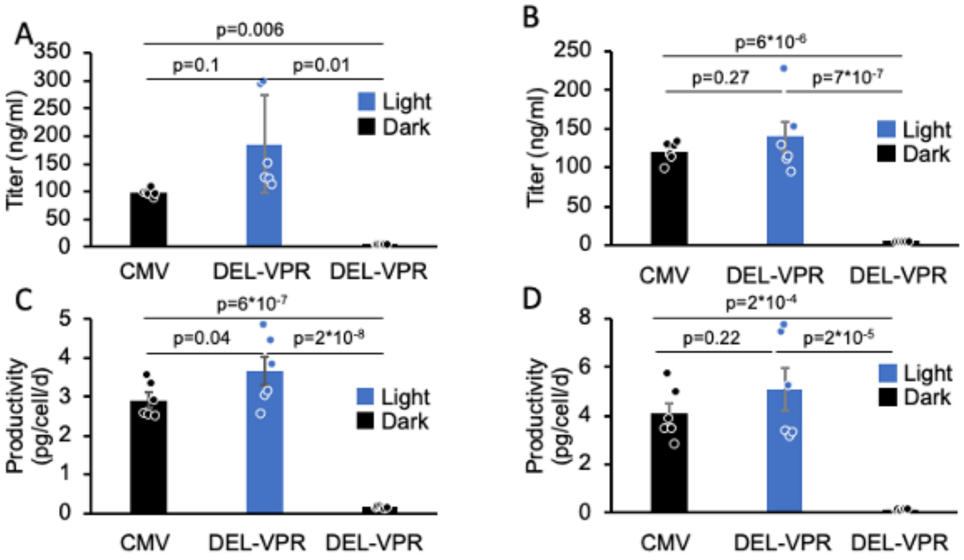


**Supplementary Figure 3: Light-dependent titration of mAb expression in HEK293T and CHO-K1.**

A) and B) show the titer of mAb of the light-induced (DEL-VPR light; 1500 µW/cm$^2$), non-induced (DEL-VPR dark), or the constitutive (CMV) conditions after 24 h expression in HEK293T or CHO-K1, respectively, quantified by ELISA. H) and I) show the productivity of mAb production of the experiment presented in C-D) in HEK293T and CHO-K1, respectively. A-D) Means of biological independent samples ± STD are presented as bars, dots indicate individual values of each sample. Statistical analysis was performed using one-way ANOVA with the Holm-Sidak post-hoc test. Plots contain six (n=6) independent samples.

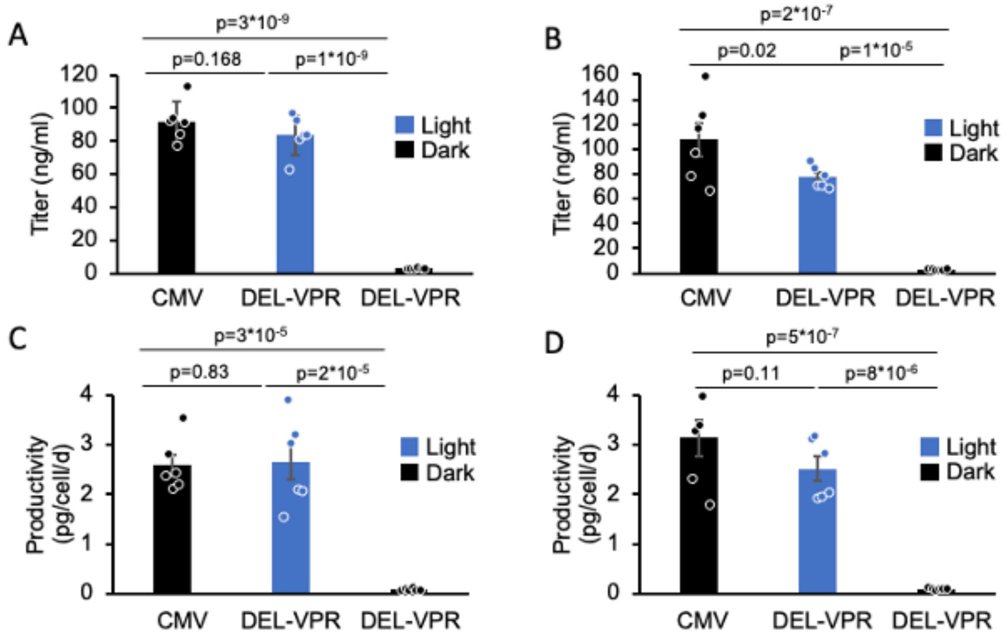


**Supplementary Figure 4: Light-dependent titration of bsAb expression in HEK293T and CHO-K1.**

A) and B) show the titer of bsAb of the light-induced (DEL-VPR; 1500 μW/cm$^2$), non-induced (DEL-VPR) or the constitutive (CMV) conditions after 24 h expression in HEK293T or CHO-K1, respectively, quantified by ELISA. C) and D) show the productivity of bsAb production of the experiment presented in A-B) in HEK293T and CHO-K1, respectively. A-D) Means of biological independent samples ± STD are presented as bars, dots indicate individual values of each sample. Statistical analysis was performed using one-way ANOVA with the Holm-Sidak post-hoc test. Plots contain six (n=6) independent samples.
