## Supplementary Table 1 for "Potent photoswitch for expression of biotherapeutics in mammalian cells by light"

List of primers used:

| Plasmid | Vector | Insert | Primer FW | Primer RV |
| --- | --- | --- | --- | --- |
| 5xC120-minP-mCherry | pcDNA3.1 | mCherry | ggtggaattcgCCACCATGGTGAGCAAGGGC | cttTCTAGATTACTTGTACAGCTCGTCC |
| 5xC120-minP-LC(mAb) | 5xC120-minP-FLuc | LC (mAb) | ttttttGGCGCGCCTAATATTGCCACCATGTTGCCATCACAACTCATTGGGTTTCTGC | aaaaaaCGCCGGCGTACTAGTCTAACACTCTCCCCTGTTGAAGCTCTTTGTGACG |
| 5xC120-minP-HC(mAb) | 5xC120-minP-FLuc | HC (mAb) | ttttttGGCGCGCCTAATATTGCCACCATGGACTGGACCTGGAGGATCCTCTTCT | aaaaaaCGCCGGCGTACTAGTTCATTTACCCGGAGACAGGGAGAGGCT |
| CMVp-LC(mAb) | pcDNA3.1 | LC (mAb) | ttttttCTCGAGACATTGATTATTGACTAGTTATTAATAGTAATCAATTACGGGGTC | aaaaaaGGCGCGCCGATCTGACGGTTCACTAAACGAGCTCTGC |
| CMVp-HC(mAb) | pcDNA3.1 | HC (mAb) | ttttttCAATTGACATTGATTATTGACTAGTTATTAATAGTAATCAATTACGGGGTC | aaaaaaGGCGCGCCGATCTGACGGTTCACTAAACGAGCTCTGC |
